# A mosquito survey along a transect of urbanization in Dschang, West Region of Cameroon, reveals potential risk of arbovirus spillovers

**DOI:** 10.1101/763755

**Authors:** Mayi Marie Paul Audrey, Bamou Roland, Djiappi-Tchamen Borel, Djojo-Tachegoum Carelle, Fontaine Albin, Antonio-Nkondjio Christophe, Tchuinkam Timoleon

## Abstract

**Background:** With the advance of globalization and the decline of wild habitats, mosquito-borne viruses are no longer confined to their original sylvatic environments and are emerging or remerging worldwide. However, little is known about the mosquito species implicated in the spillovers of these viruses from their enzootic cycles.

**Method:** We conducted an entomological field survey to catalogue the relative abundance of the *Culicidae* fauna in rural, peri-urban and urban areas in the Dschang locality in West Cameroon. Mosquitoes were collected from March-April and July-August 2019 at both aquatic and adult stages, and identified using stereomicroscopes and morphological identification keys.

**Results:** A total of 1,401 mosquitoes belonging to 4 genera and 26 species were collected (n=427, 470 and 504 in rural, peri-urban and urban areas respectively). The most abundant species *Aedes africanus* (45.47%) as well as *Culex moucheti* (8.92 %) were encountered in peri-urban and rural environments. Species like *Culex duttoni* (14.49%)*, Culex quinquefasciatus* (2.35%) and *Aedes aegypti* (1.36%) were solely found in urban area. *Aedes albopictus* (14.42%)*, Culex (Culiciomyia)* (6.57%), *Culex tigripes* (1.43%) and *Eretmapodites chrysogaster* (0.64%) on the other hand were collected in the three study sites. Importantly, all these species have been suspected or incriminated as vectors of many arboviruses.

**Conclusion:** This study identified potential sylvatic, urban and bridge-vectors that can play a role in current or future virus emergence in Cameroon. Further investigations are needed to assess if arboviruses are circulating in these areas and to study the vector role of each mosquito species in arbovirus transmission.

## Background

Mosquito-borne viruses are becoming a serious public health problem worldwide as they spill over from their wild sylvatic habitat to emerge in many urban areas in all continents wherever competent vectors are present. Most of the major mosquito-borne viruses of medical importance that cause disease outbreaks in human populations without the requirement of amplification by a secondary non-human vertebrate host originated from the African continent (like Yellow fever virus, Chikungunya virus and Zika virus).

In a recent review, Braack and colleagues [1] listed 36 major arthropod-borne viruses (arboviruses) indigenous to Africa with their respective geographic distribution and known mosquito vector species. In Cameroon, the authors recorded many arboviruses, which include Dengue virus (DENV), Ntaya virus (NTAV), Spondweni virus (SPOV), Yaounde virus (YAOV) and Yellow fever virus (YFV) from the *Flavivirus* genus, *Flaviviridae* family; Chikungunya virus (CHIKV), Semliki Forest virus (SFV) and Sindbis virus (SINV) from the *Alphavirus* genus, *Togaviridae* family; Rift Valley Fever virus (RVFV) from the *Phlebovirus* genus, *Bunyaviridae* family; and Bunyamwera virus (BUNV), Bwamba virus (BWAV) and Ilesha virus (ILEV) from the *Orthobunyavirus* genus, *Bunyaviridae* family. These findings originate from many studies that have investigated the prevalence of arboviruses in Cameroon mainly based on the serodiagnostic of blood samples collected from humans [2–12]. However, few studies have investigated potential urban vectors or bridge vectors responsible for spillovers of these viruses from the forest into anthropized environments through entomological surveys [2, 13, 14], and very few have searched for viruses in mosquitoes in Cameroon [15, 16].

To address this gap in knowledge, we made an up-to date catalogue of the diversity and abundance of mosquito species collected in three habitats along a transect of urbanization in Dschang, West Region of Cameroon. This survey would help to identify potential sylvatic, urban and bridge-vectors that can play a role in current or future virus spillover from wild to more urbanized areas. It is also an important initial step to assess the health risk of humans and animals living in these habitats to such diseases.

## Methods

### Description of the study sites

This study was carried out in rural (sylvatic), peri-urban and urban areas of Dschang Sub Division, within the Menoua Division. The rural and peri-urban habitats were located in Fonakeukeu (N: 05 24.48.1, E: 010 04.44.7) and Toutsang (N: 05 25.34.5, E:01004.10.7) villages respectively while the urban habitat was located in the town of Dschang (N: 05 26.805, E: 010 03.8404). They all lie in the highland area of western Cameroon that exhibits a sub-tropical climate characterized by two seasons; a dry season of 4 months (from mid-November to mid-March) and a rainy season of 8 months (mid-March to mid-November). Annual average rainfall and temperature are of 321 mm and 21.6°C respectively [10]. The urban area (Dschang) is characterized by a shrub-like vegetation, an important hydrographic network and many artificial breeding sites, ideal to maintain the urban cycle of mosquitoes and their arboviruses. The peri-urban (Toutsang) and rural (Fonakeukeu) are found at about 2km and 5 km respectively from the urban area (Dschang). Raffia palm bushes are common in these areas and are found along the valleys and streams which offer natural breeding sites that can easily maintain the sylvatic cycle of mosquitoes and their arboviruses.

### Mosquito sampling and identification

Field surveys were made in rural, peri-urban and urban areas during the rainy season from March-April and July-August 2019. Mosquitoes were collected in these areas three times per month at immature stages in available breeding sites (abandoned tires, riverbeds and floor pools) and at adult stages using sweep nets (to catch males, females and bloodfed females resting on the vegetation). Immature stages were reared to adults before identification. Morphological identification of species was done using stereomicroscopes and morphological identification keys [17–19].

## Results

A total of 1,401 mosquitoes belonging to 4 genera and 26 species were collected in the three areas (n=427 in the rural area, n= 470 in the peri-urban area and n=504 in the urban area) (Table 1). Out of the 26 species, *Aedes africanus* (n=637, 45.47%) were the most abundant, followed by *Culex duttoni* (n=203, 14.49%), *Aedes albopictus* (n=202, 14.42%), *Culex moucheti* (n=125, 8.92%) and *Culex (Culiciomyia)* (n=92, 6.57%) (Table 1). Interestingly, *Aedes africanus* and *Culex moucheti* were only found in rural (Fonakeukeu) and peri urban (Toutsang) areas while *Culex duttoni, Culex quinquefasciatus and Aedes aegypti* were only found in urban area (Dschang). Species from the *Culex Culiciomyia* group, *Eretmapodites chrysogaster* group, *Culex tigripes* and *Aedes albopictus* on the other hand were caught in the three sites (rural, peri urban and urban). More importantly, all these species have been suspected or incriminated as vectors of many arboviruses (Table 1).

**Table 1:**
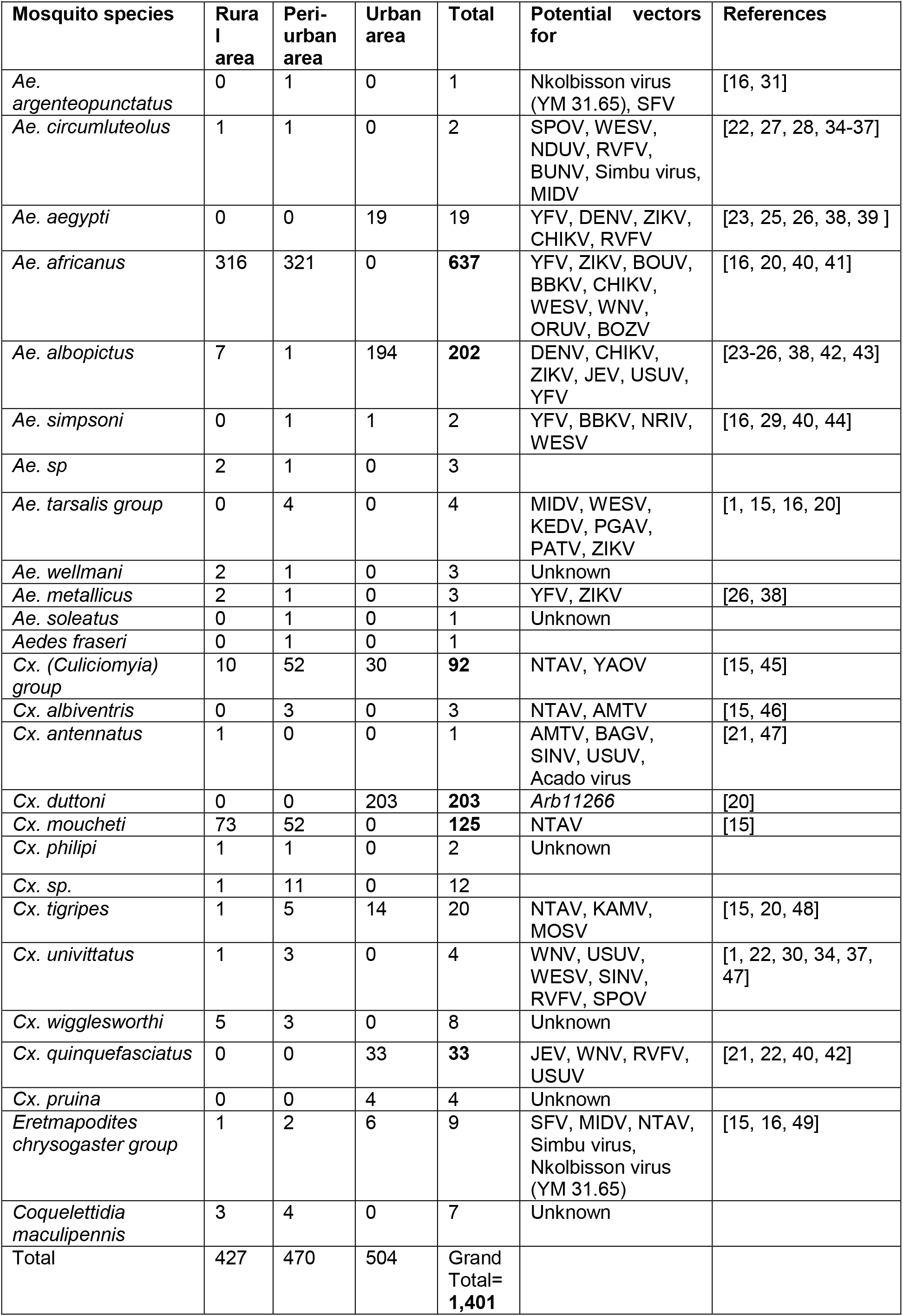
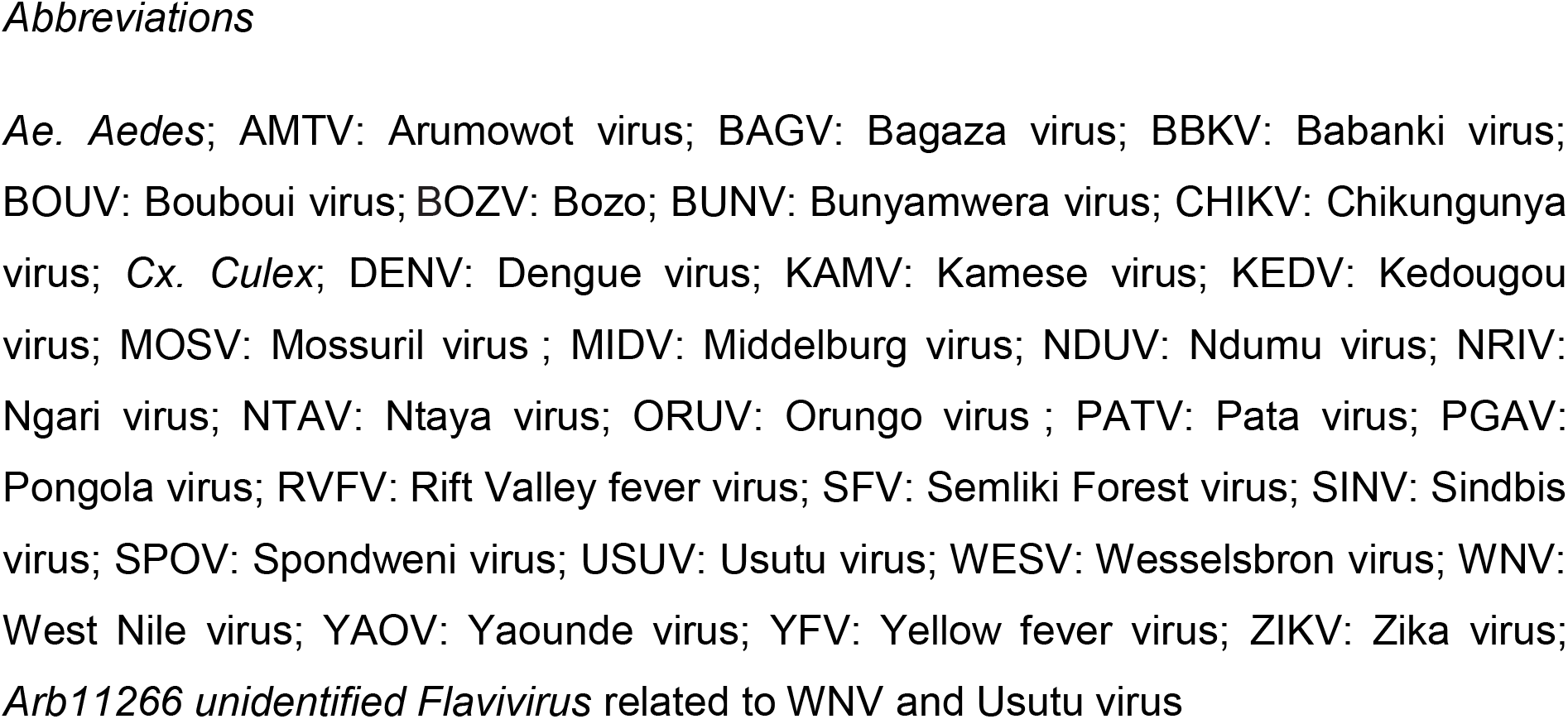
Mosquito species collected in Dschang and potential vectors of arboviruses worldwide

## Discussion

This work aimed to catalogue the *Culicidae* fauna in urban, peri-urban and rural areas of Dschang, West Cameroon, that can be implicated as sylvatic, urban or bridge vectors in virus transmission. A high diversity of mosquito species was found in the locality with about 65.38% of the total mosquitoes collected identified as potential vectors of arboviruses (Table 1).

*Aedes africanus* was the most abundant mosquito species in rural and peri-urban areas. This species can transmit several arboviruses and has been incriminated as vector of YFV and CHIKV [20]. *Culex duttoni*, the second most abundant mosquito species found in the urban area, has been found to carry arboviruses like *Arb11266* [20]. *Culex quinquefasciatus*, another species found exclusively in the urban area, has been reported to transmit West Nile Fever (WNV) [21] and RVFV [22]. *Aedes albopictus* is known vector of CHIKV [23] and has been suspected in Zika virus transmission in Gabon [24]. *Culex Culiciomyia* species and *Culex moucheti* were implicated in the transmission of NTAV [15].

Other mosquito species although collected in small numbers in our study area, have also been identified as vectors of many arboviruses. These are; *Aedes aegypti*, vectors of DENV, YFV, CHIKV [23, 25, 26]; *Aedes circumluteolus*, vectors of SPOV, RVFV, BUNV [27, 28]; *Aedes metallicus*, vectors of YFV [26]; *Aedes simpsoni*, vectors of YFV [29]; *Culex univittatus*, vectors of SINV [30]; *Aedes argenteopunctatus*, vectors of Nkolbisson virus (YM 31.65) and SFV [16, 31]. Interestingly, a high aggressiveness of mosquitoes was noted during sampling and few of the collected bloodfed mosquitoes (like *Eretmapodites chrysogaster* group, *Cx. moucheti* and *Ae.* a*fricanus*) were found with human blood (Unpublished data).

Arboviruses that have been found circulating in Cameroon include DENV, YFV, CHIKV, Zika virus (ZIKV), ONNV (Onyong Onyong virus), NTAV, SPOV, Wesselsbron virus (WESV), SFV, SINV, Middelburg virus (MIDV), WNV, Tahyna virus (TAH), BUNV, Uganda S virus (UGSV), YAOV, RVFV, BWAV and ILEV [1, 2, 4, 6, 8, 9, 12]. However, few mosquitoes have been incriminated as potential vectors of these arboviruses in Cameroon. These few include; *Aedes tarsalis* group (viruses isolated: MIDV, WESV), *Aedes argenteopunctatus* (virus isolated: Nkolbisson (YM 31.65)), *Aedes africanus* (virus isolated: WESV), *Aedes simpsoni* (virus isolated: WESV), *Culex albiventris* (virus isolated: NTAV), *Culex Culiciomyia* (virus isolated: NTAV), *Culex moucheti* (virus isolated: NTAV), *Culex tigripes* (virus isolated: NTAV) and *Eretmapodites chrysogaster* group (viruses isolated: MIDV, Simbu virus) [15, 16].

Many mosquito species sampled in this study, such as *Aedes africanus*, *Aedes aegypti*, *Aedes tarsalis*, *Aedes albopictus* and *Aedes metallicus*, have been reported to be potential vectors of ZIKV and DENV in previous studies. Yet, ZIKV has been detected in Cameroon in Yaoundé [5, 9] and in the Fako Division [4]. Furthermore, a recent study found 11.7% and 10.4% of DENV circulating in Dschang and Bagangté respectively [10], all situated in the western highland. Another study revealed acute dengue in children living in urban and semi-urban areas of Dschang [32]. A mosquito-based arbovirus surveillance system might therefore help as a supplement to assess the presence of such arboviruses in the locality and in the whole country.

## Conclusion

This study provides an up-to date catalogue of the *Culicidae* fauna along a transect of urbanization and identify the potential mosquito vectors that may be involved in arbovirus transmission in West Cameroon. Based on these preliminary findings, in addition to the recent detection of DENV in the locality, further researches are required to assess the prevalence of arboviruses in these areas and to study the vector role of these mosquito species in arbovirus transmission as well as to assess their anthropophily by blood meal source identification. A surveillance system based on the detection of pathogen genomic material in mosquito excreta has been recently developed [33]. This system is time and cost effective as compared to the standard method that relies on processing thousands of individual mosquitoes and would offer the opportunity to screen virus transmission in wider and wilder areas.

## Authors’ contributions

Mayi Marie Paul Audrey, Bamou Roland, Djiappi-Tchamen Borel and Tchuinkam Timoleon, designed the study. Mayi Marie Paul Audrey, Bamou Roland, Djiappi-Tchamen Borel and Djojo-Tachegoum Carelle collected data in the field and analyzed themin the laboratory. All the authors drafted and revised the article for scientific and intellectual content. All the authors read and approved the final manuscript

